# Comparative analysis of the genomes of *Stylophora pistillata* and *Acropora digitifera* provides evidence for extensive differences between species of corals

**DOI:** 10.1101/197830

**Authors:** Christian R Voolstra, Yong Li, Yi Jin Liew, Sebastian Baumgarten, Didier Zoccola, Jean-François Flot, Sylvie Tambutte, Denis Allemand, Manuel Aranda

## Abstract

Stony corals form the foundation of coral reef ecosystems. Their phylogeny is characterized by a deep evolutionary divergence that separates corals into a robust and complex clade dating back to at least 245 mya. However, the genomic consequences and clade-specific evolution remain unexplored. In this study we have produced the genome of a robust coral, *Stylophora pistillata*, and compared it to the available genome of a complex coral, *Acropora digitifera*. We conducted a fine-scale gene-based analysis focusing on ortholog groups. Among the core set of conserved proteins, we found an emphasis on processes related to the cnidarian-dinoflagellate symbiosis. Similarly, genes associated with the algal symbiosis were also independently expanded in both species, but both corals diverged on the identity of ortholog groups expanded, and we found uneven expansions in genes associated with innate immunity and stress response. Our analyses demonstrate that coral genomes can be surprisingly disparate. Importantly, if the patterns elucidated here are representative of differences between corals from the robust and complex clade, the ability of a coral to respond to climate change may be dependent on its clade association.

## Introduction

Coral reefs are ecologically and economically highly important marine ecosystems, as they provide biodiversity hotspots for a large diversity of species and serve as a food source for millions of people ^1,2^. Despite their importance, coral reefs are threatened by a combination of local (e.g., overfishing, eutrophication, pollution) and global (e.g., ocean warming and ocean acidification) factors that cause an increase of coral disease and coral bleaching, which in many cases lead to the ultimate death of affected coral colonies ^3-5^. Over the last decades, coral reef cover was significantly decimated and one-third of reef-building corals face elevated extinction risk from climate change and local impacts ^6^. For this reason, it is important to understand the factors that contribute to ecosystem resilience.

At the heart of these ecosystems are the so-called coral holobionts, which provide the foundation species of reefs and consist of the coral animal host, its endosymbiotic photosynthetic algae, and a specific consortium of bacteria (among other organisms) ^7,8^. While recent research highlights the contribution of all holobiont compartments to coral resilience ^9-13^, the majority of studies focus on the diversity of algal and bacterial symbionts associated with corals or on gene expression of the host under an array of stressors or across different environments ^14-21^. Hence, although coral species display differing sensitivities to environmental stress ^9^, the genomic underpinnings of coral resilience are not clear.

A recent study by Bhattacharya, et al. ^22^ conducted a comparative analysis incorporating genomic and transcriptomic data from 20 coral species. Focusing on the orthologs conserved across all analyzed corals, the authors describe the presence of a variety of stress-related pathways (e.g., apoptotic pathways, reactive oxygen species scavenging pathways, etc.) that affect the ability of corals to respond to environmental stress. Importantly, the authors could show that corals harbor a highly adaptive gene inventory where important genes arose through horizontal gene transfer or went through rounds of evolutionary diversification. Similarly, the recently published genome of *Acropora digitifera* ^23^ highlights that the innate immunity repertoire of corals is presumably and notably more complex than those of the cnidarians *Nematostella vectensis* ^24^ and *Hydra magnipapillata* ^25^. Seemingly so, the innate immunity repertoire is also more complex than that of the symbiotic anthozoan sea anemone *Aiptasia* ^26^. This has potential implications for our understanding of coral responses to environmental change. Unfortunately, it is not straightforward to determine what a ‘typical’ coral genome looks like. This is because the phylogeny of scleractinian corals is characterized by a deep evolutionary split that separates corals into a robust and complex clade dating back to at least 245 mya ^27,28^. Hence, several important questions, such as how well the available genome of *Acropora digitifera* indeed reflects general coral-specific traits and to what extent species from both coral clades diverged since their separation (giving rise to different adaptations) are currently unanswered due to the dearth of coral genomes. To this end, the Reef Future Genomics (ReFuGe) 2020 consortium has formed to sequence 10 hologenomes of coral species representing different stress susceptibilities in order to better understand conserved and lineage-specific traits, but a comprehensive analysis is pending ^9^.

In this study, we produced and analyzed the genome of *Stylophora pistillata*, a representative of the robust clade of corals, and compared it to the available genome of the complex coral *A. digitifera*. We were specifically interested in a comparison of (1) the set of orthologous genes, (2) species-specific genes, and (3) genes that were independently expanded in either of the genomes or both. These three classes of genes, we reasoned, provide complementary insight into the evolutionary history of both corals, and may highlight important species-specific adaptive processes ^29,30^. Further, such a comparative analysis may pinpoint genomic differences that arose from the different evolutionary trajectories that occurred in coral species from either clade and, as such, may represent clade-specific differences.

## Results

### Genome size and genic composition

We assembled 400 Mb of the genome of the coral *S. pistillata* (Table 1, Fig. S1) with a scaffold N50 of 457 kb, representing ~92% of the 434 Mb genome as estimated via FACS (Fig. S2). 358Mb were assembled into contigs, with a contig N50 of about 24 kb (Table 1, Table S1). We identified 25,769 protein-coding genes encoded in the *S. pistillata* genome, of which 89% retrieved functional annotation from protein databases (Table 1, Table S2). The genome size and the number of genes are comparable to the draft genome of *A. digitifera* that features a total scaffold length of about 419 Mb with a scaffold N50 of 191 kb and 23,523 protein-coding genes (Table 1). However, genome completeness as assessed by CEGMA ^31^ was considerably higher in *S. pistillata* with about 94.76% of the core eukaryotic genes present compared to 82.26% in *A. digitifera* (Table S3).

**Table 1.** Assembly statistics for the genomes of *S. pistillata* and *A. digitifera*.

|  |  | *Stylophora*<br>*pistillata* | *Acropora*<br>*digitifera* |
| --- | --- | --- | --- |
| **Genome** | Genome file used | v1.0 | v1.0 (Jul 2011) |
|  | Estimated genome size (Mb) | 434 | 420 |
|  | Total scaffold length (bp) | 400,108,361 | 419,317,576 |
|  | Scaffold N50 (bp) | 457,453 | 191,489 |
|  | Total contig length (bp) | 358,078,850 | 364,965,673* |
|  | Contig N50 (bp) | 24,388 | 10,700* |
|  | GC content, N excluded (%) | 38.5 | 39.0 |
| **Genes** | Number of genes (longest transcript per locus) | 25,769 | 23,523 |
|  | Mean gene length (bp) | 8,432 | N/A |
|  | Gene model EST support (%) | 82.1 | 78* |
| **Exons** | Mean coding region length (bp) | 2,086 | 1,707* |
|  | Number of exons per gene | 7.9 | 7.0* |
|  | Mean length (bp) | 266 | 230 |
|  | Total length (Mb) | 53.8 | 40.2 |
| **Introns** | Genes with introns (%) | 96.6 | N/A |
|  | Mean length (bp) | 918.3 | N/A |
|  | Total length (Mb) | 162.1 | N/A |
| **Intergenic** | Average length (bp) | 6,333 | N/A |

\*: from Shinzato et al., 2011.
Some statistics were not available as the *A. digitifera* v1.0 gff3 file was not made public.

To obtain general insight into the genic composition of coral genomes, we performed a BLASTP search with the gene sets encoded in both genomes against the ‘nr’ protein database (see Materials & Methods). The vast majority of genes from both species had best matches to *Aiptasia* (48.36% for *S. pistillata* vs. 43.82% for *A. digitifera*) and *Nematostella* (23.54% for *S. pistillata* vs. 25.82% *A. digitifera*) (Fig. 1a). The remaining genes generally matched non-cnidarian proteins or had no matches (7.30% *S. pistillata* vs. 10.55% for *A. digitifera*), presumably representing lineage-specific or species-specific genes. Strikingly, when this analysis was extended to allow for inter-coral matching, pronounced differences were revealed between both coral species (Fig. 1b). In particular, we found that matches of *A. digitifera* genes to *S. pistillata* genes were highly disproportional (17,866 *A. digitifera* genes matched to 10,945 *S. pistillata* genes, *p* < 10^-300^, Fisher’s exact test), indicating potential pervasive gene duplication in *A. digitifera*. In addition, *A. digitifera* exhibited significantly fewer matches to the anemones *Aiptasia* (1,942 genes) and *Nematostella* (1,011 genes) than *S. pistillata* (6,994 gene matches to *Aiptasia*, 3,437 gene matches to *Nematostella*) (Fisher’s exact test, *p* < 10^-300^ and *p* < 10^-283^, respectively), pointing towards increased divergence of protein sequences in *A. digitifera*.

**Figure 1.**
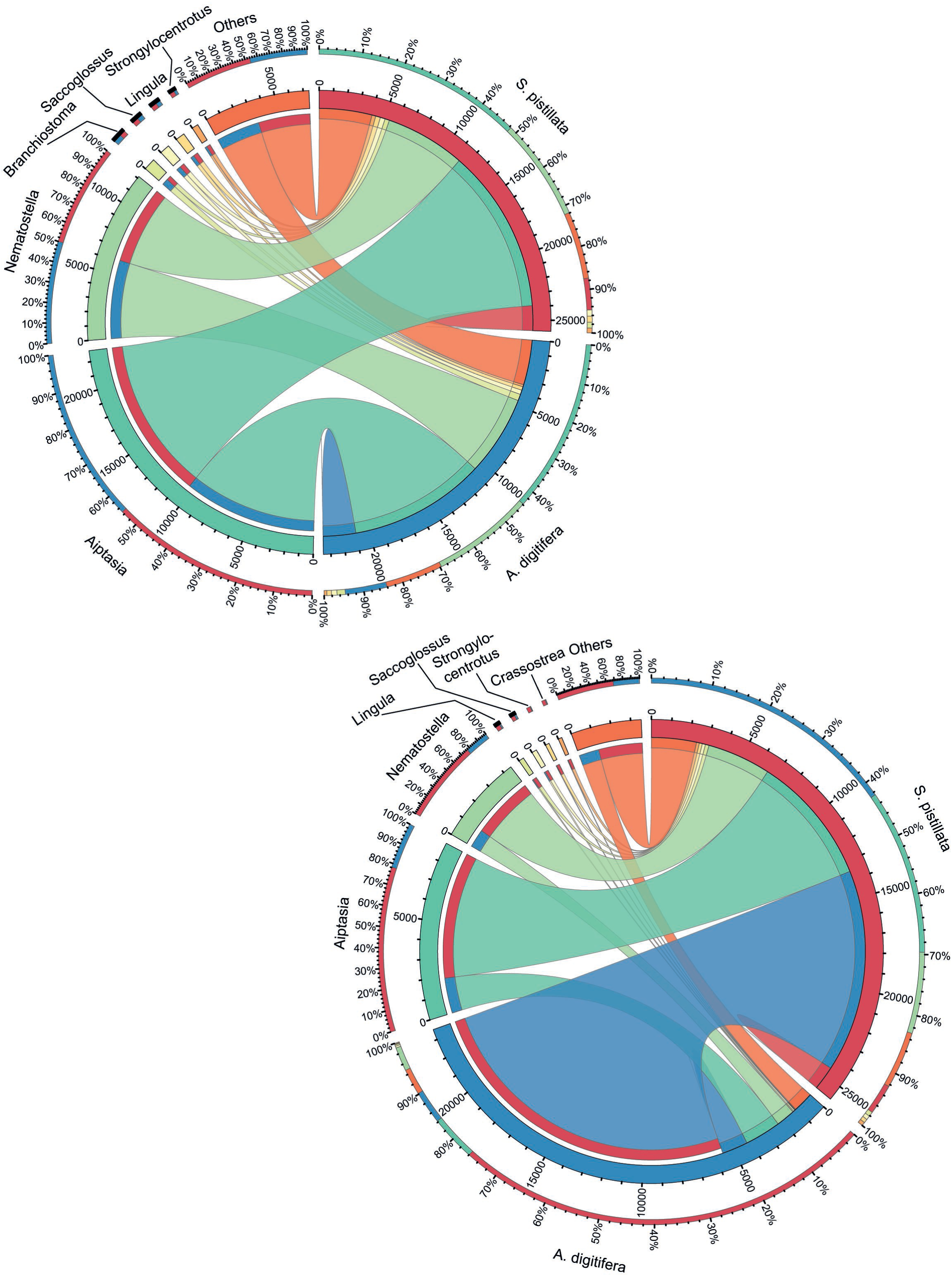
Chord diagrams showing genomic gene compositions of *S. pistillata* and *A. digitifera*. For both diagrams, best matches to both corals and top 6 genera are shown.(A) Both coral genomes appear similar in composition when queried against non-coral sequences. As expected, most of the matches were to other cnidarian species, such as *Aiptasia* or *Nematostella*. (B) If coral genomic gene sets are allowed to match against each other, many more *A. digitifera* genes match homologs in *S. pistillata* homologs than *vice versa*. As a consequence of the asymmetrical matching, the number of *A. digitifera* matches to other genera is vastly reduced.

### Conservation of protein-encoding genes

We first compared the genomes at the protein level, considering 25,769 *Stylophora* and 23,523 *Acropora* protein-encoding genes. The proteins were classified into four categories according to their evolutionary relationships (Fig. 2). The first category includes *Stylophora* proteins with one clearly identifiable counterpart in *Acropora* and *vice versa* (one-to-one orthologs). The function of these proteins is likely conserved and can be interrogated to infer core functions of coral genomes. This approach was employed in a recent study ^22^, where the authors collated and queried data from 20 coral species (including *S. pistillata* and *A. digitifera*) to elucidate four major issues in coral evolution, i.e. coral calcification, environmental sensing, symbiosis machinery, and the role of horizontal gene transfer (HGT). Here, we used reciprocal best matches that produced 6,302 protein pairs classified as one-to-one orthologs (24% of *S. pistillata* and 27% of *A. digitifera* proteins) (Supplementary Dataset S1). The second category included proteins in which gene duplication has occurred in one or both species after divergence, resulting in “many-to-one” and “many-to-many” ortholog relationships, respectively. This group consisted of 2,747 *S. pistillata* and 2,900 *A. digitifera* proteins (11% of *S. pistillata* and 12% of *A. digitifera* proteins) that presumably harbor genes that expanded independently in both lineages (Supplementary Dataset S2). We hypothesize that the presence of species-specific gene expansions likely reflects functions relevant to either species- or clade-specific evolution. The third category included 15,442 *S. pistillata* and 12,925 *A. digitifera* proteins (60% and 56%, respectively) that have homologs in corals or other species, but without easily discernable orthologous relationships between corals. The high number of this group of proteins likely also reflects our conservative approach for ortholog identification (see Materials & Methods). Finally, the fourth group consisted of 1,278 *Stylophora* and 1,396 *Acropora* proteins that have no detectable homologs in any other species. These proteins putatively belong to the class of taxonomically restricted genes (TRGs) that might be encoded by lineage-specific or fast evolving genes ^29^.

**Figure 2.**
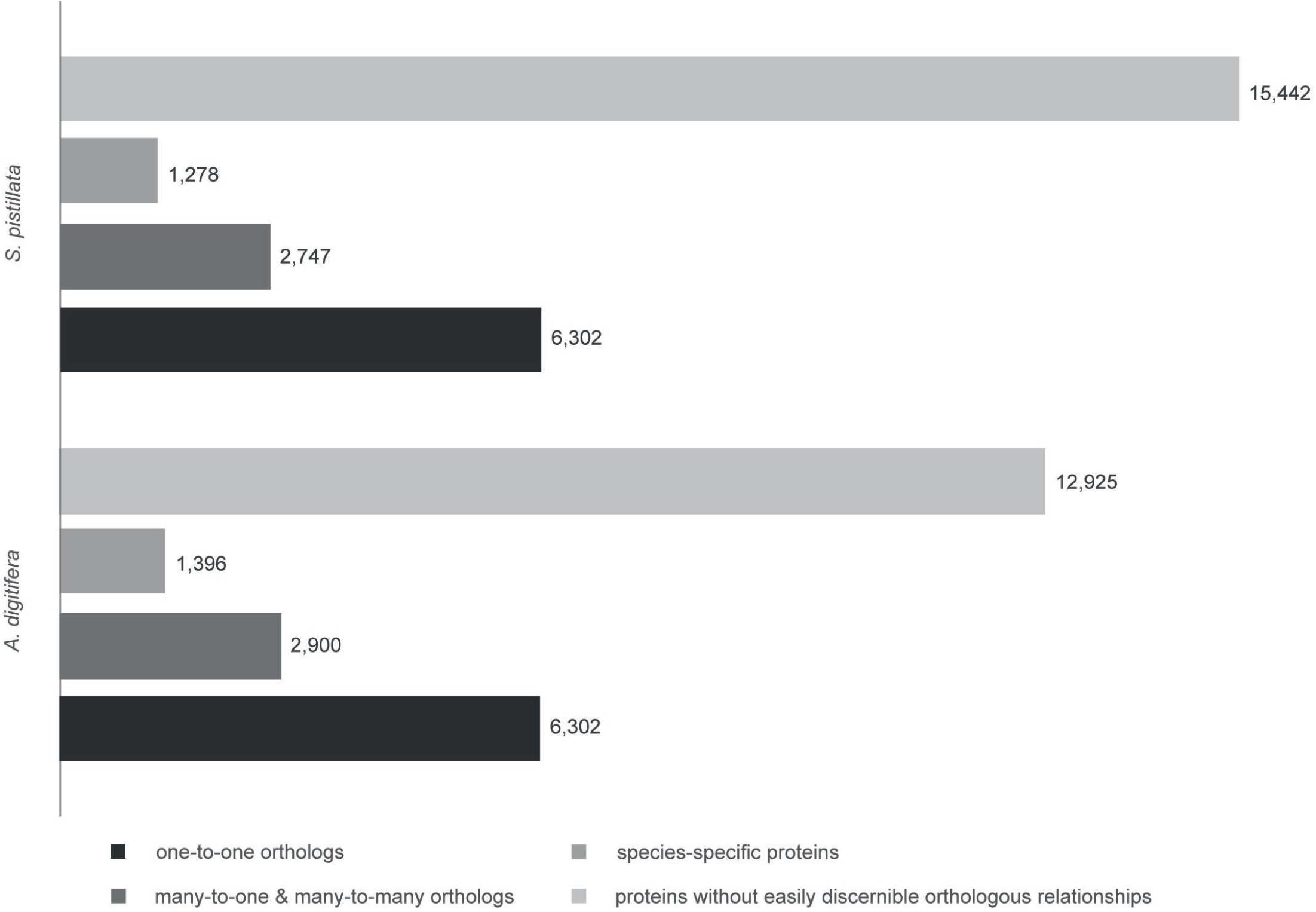
Classification of genomic protein sets of *S. pistillata* and *A. digitifera* according to evolutionary relationship. 25,769 *S. pistillata* proteins were compared to 23,523 *A. digitifera* proteins and assigned to four categories: (i) one-to-one orthologs (blue), (ii) many-to-one and many-to-many orthologs (green), (iii) proteins without easily discernible orthologous relationships (teal), and (iv) species-specific proteins without homologs in other species (dark blue).

### The core set of conserved proteins highlight processes relevant to coral evolution

The average sequence identity of the one-to-one orthologs of *S. pistillata* and *A. digitifera*, which are presumably at least ~245 mya apart ^27^ was 62% on the protein level. By comparison, average sequence identity between *Anopheles* and *Drosophila*, which are separated by approximately the same time ^32^ was estimated to be 56% in a previous study ^30^. This indicates that despite the comparable divergence time in both comparisons, coral proteins diverge at a lower rate than insect proteins, possibly because corals have much longer generation times ^9^, although a substantial portion of the genome can evolve at elevated rates ^33^. At the same time, the average sequence identity between orthologs shared by humans and pufferfish is 61%, and these species are approximately 450 million years apart ^34^, indicating that corals are not at the lowest end of divergence rates.

The one-to-one orthologs constitute a core of conserved functions that encode for basic biological processes, which is corroborated by a Gene Ontology (GO) based analysis (Supplementary Dataset S3). The 50 most common GO terms are associated with regulation of metabolism and cellular processes, organelle function, and notably, nitrogen-related metabolic processes, the regulation of which were previously discussed as central to coral holobiont functioning ^35^.

To test whether ortholog groups across a range of sequence similarities were enriched for certain biological processes, we divided the set of one-to-one orthologs into three groups: orthologs displaying ≥ 50%, between < 50% and ≥ 30%, and those displaying < 30% sequence similarity and tested for Gene Ontology enrichment (Supplementary Dataset S4). The group of highly conserved orthologs was, as expected, enriched for genes associated with housekeeping processes, such as transcription, translation, and ribosomes. In comparison, the group displaying similarity between 50% and 30% were enriched for orthologs associated with endocytosis, immune system activation (NFkB), and superoxide metabolic processes, which putatively play a role in the endosymbiosis. In contrast, the group of orthologs with similarity < 30% was enriched for processes playing a role in cell adhesion, cell junctions, and calcium ion binding. In this regard, it is interesting to note that a recent study ^36^ found an unexpected diversity of structural components of septate junctions in cnidarians and suggested that genes involved in the formation of septate junctions may determine coral resistance to ocean acidification.

### Gene expansions and reductions point to a set of common and species-specific processes related to cnidarian-dinoflagellate endosymbiosis

A functional enrichment analysis using Gene Ontology information highlighted several biological processes that were enriched in the category of orthologs with “many-to-one” and “many-to-many” relationships, and several of these processes were shared between both coral species, although the majority of enriched processes were species-specific (Supplementary Dataset S5). The common processes included several immunity-related GO categories associated with the regulation of NFkB and in particular interferon production, but also categories related to cell adhesion and bicarbonate transport. Arguably, all these processes are related to the cnidarian-dinoflagellate endosymbiosis. Immunity-related GO terms were also present in the enriched categories specific to *S. pistillata*, but other GO terms prevailed. For instance, several processes related to receptor-mediated endocytosis, amine metabolism, osmosis, apoptosis, and hyperoxia were enriched. Again, these processes are conceivably related to the symbiotic relationship with zooxanthellae. In *A. digitifera* we also found enrichment of processes related to innate immunity, in particular of genes associated with the Toll signaling pathway, interleukin production, and bacterial detection. Other enriched processes specific to *A. digitifera* were notably different however, such as miRNA metabolism, cytoskeletal organization, hydrogen peroxide metabolism, proteolysis, and pyroptosis.

### Uneven expansions of proteins related to the immune system

The group of proteins with gene duplications in one or both species revealed many uneven expansions (or reductions) as highlighted by the observation that in many cases a single protein in either species had multiple counterparts in the other species. Genes experiencing multiple rounds of duplications in either one or both species are arguably among the most interesting proteins to look at, as they may reveal information on processes independently selected in either or both species. For this reason, we further looked into ortholog groups with at least 3 proteins in either one or both corals.

Both coral species expanded genes related to innate immunity receptors. For instance, we discovered 3 cases where a gene encoding for a NOD-like receptor family member (NLRC3) was independently expanded in both corals (Supplementary Dataset S6). Further, we found 1 case where a gene encoding for a TLR (Toll-like receptor 1), and another case where a gene encoding for a TNFR (Tumor necrosis factor receptor superfamily member 1) showed independent duplication in both corals (Supplementary Dataset S6). Importantly, the ortholog groups showed different degrees of expansions in both corals. In three of the above cases, we found more duplicated genes in *A. digitifera* than in *S. pistillata*. In contrast, in only one of the above cases we found more genes in *S. pistillata* than *A. digitifera* (Supplementary Dataset S6). We found an extreme case of expansion for one of the NOD-like receptor family members, where *A. digitifera* harbored 52 proteins in comparison to *S. pistillata* that harbored only 5 proteins (Fig. 3, Supplementary Dataset S6). Overall, *A. digitifera* had a stronger tendency to show ‘extreme’ expansions (10 ortholog groups with more than 10 proteins) than *S. pistillata* (3 ortholog groups with more than 10 proteins) (Supplementary Dataset S6). Besides innate immunity receptors, we also found 2 ortholog groups related to biomineralization, i.e. homologs of Carbonic Anhydrase (CA) and Bone Morphogenetic Protein 1 (BMP-1), to be expanded in both species, but at very similar levels (3 vs. 4 proteins for CA and 3 proteins each for BMP-1 for *A. digitifera* and *S. pistillata*, respectively) (Supplementary Dataset S6).

**Figure 3.**
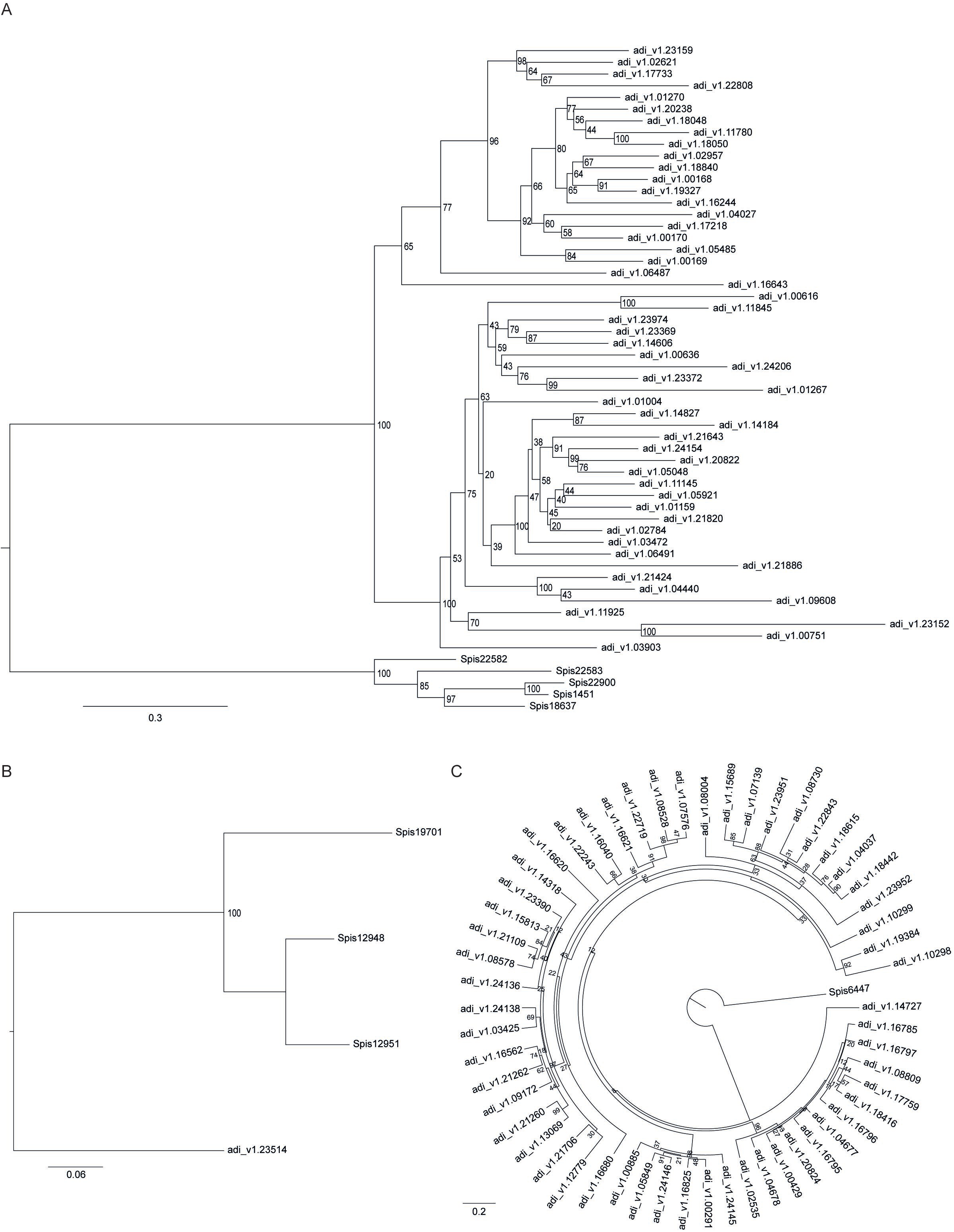
Gene expansion of orthologs in *S. pistillata* and *A. digitifera*. (A) Ortholog expansion displaying a many-to-many relationship for a NOD-like receptor family members, in which *A. digitifera* harbors 52 proteins in comparison to *S. pistillata* that harbors only 5 proteins. (B) Ortholog expansion displaying a many-to-one relationship of a TRAF (TNF receptor-associated factor 3) homolog with expansion in *S. pistillata*. (C) Ortholog expansion displaying a many-to-one relationship of a member of the NOD-like receptor family member (NLRC3) with a particular pronounced expansion in *A. digitifera* (55 genes) and only one corresponding counterpart in *S. pistillata.*

For the groups of proteins with gene duplications in only of the coral species (Supplementary Dataset S7), we identified 167 genes in *S. pistillata* that mapped to groups of three or more genes in *A. digitifera*. Most notably, a member of the NOD-like receptor family member (NLRC3) gave rise to 55 genes in *A. digitifera* with 1 counterpart in *S. pistillata* (Fig. 3). Similarly, 191 genes from *A. digitifera* mapped to groups of 3 or more genes in *S. pistillata*. Among these, we found homologs of innate immunity related proteins, namely TRAFs (TNF receptor-associated factor 3) (Fig. 3) and TLRs (Toll-like receptor 2) to be present with 3 copies, and a homolog of peroxidasin, important for oxidation-reduction, to be present with 5 copies in *S. pistillata*.

### Discussion

The inference of evolutionary relationships within the Scleractinia is an ongoing subject of debate, complicated by the phenotypic plasticity in skeletal growth forms and unusual slow mitochondrial DNA sequence evolution ^37,38^. Nevertheless, a major distinction into two clades ("complex" and "robust" corals) within the Scleractinia dating back to about 245 mya ^27^ is corroborated by molecular analyses ^37,39,40^. In line with this estimate, scleractinian corals first appeared in the fossil record about 245 mya ^41^. However, the genomic consequences of this deep divergence remain unexplored. In this study we have assembled the genome of the robust coral *S. pistillata* and compared it to the available genome of the complex coral *A. digitifera* to gain a first look at coral species differences from both clades on a genomic scale.

To thoroughly understand the extent of conservation at the protein level, we followed an ortholog-based approach where we assigned proteins into four categories according to their evolutionary relationships: (i) one-to-one orthologs, (ii) many-to-one and many-to-many orthologs, (iii) proteins without easily discernible orthologous relationships, and (iv) species-specific proteins without homologs in other species. Notably more than half of the proteins from both species could not be assigned clear orthologous relationships, putatively indicating the substantial divergence associated with the deep evolutionary split between both corals. This is further corroborated by the genic composition results, which shows that more than two thirds of proteins in both species match to homologs in other cnidarian species (*Aiptasia* and *N. vectensis*).

Of the remaining other half of proteins from the ortholog-based analysis, about two thirds of the proteins (6,302 protein pairs) displayed clear one-to-one relationships, which can be considered core proteome members. This number closely resembles the number of one-to-one orthologs identified by Debashish *et al*. ^22^ in a large metaanalysis of available coral genomes and transcriptomes (4,751 ortholog pairs). Within the set of conserved orthologs across corals, the authors characterized sets of proteins responsible for biomineralization, environmental sensing, and response to temperature, light, and pH. Our analysis of enriched GO terms largely supports the results of the study by Debashish *et al*. ^22^ and further highlights the overarching emphasis on processes related to the cnidarian-dinoflagellate symbiosis in the coral host core set of conserved proteins.

A particular interesting category of orthologs is comprised of those with many-to-one and many-to-many relationships. Family expansion and reduction can be measured in different ways. The most basic measure is to annotate proteins to their domains and compare the normalized domain count between genomes. Although this is straightforward way to assess overall similarities and differences between genomes, it does not provide information on relatedness of proteins with the same domains/domain compositions. A better resolution is provided by an analysis of the enriched functions of the many-to-one and many-to-many orthologs. About 10% of proteins from both genomes fall into this category. Although this group is less strictly defined than the group of one-to-one orthologs, they can still be assigned to a single ancestral gene, and hence, imply duplication within the species, i.e. after both species diverged. As such, analysis of these proteins allows inferences on adaptations, e.g. to different environments or life strategies. Following this reasoning, we interrogated the group of orthologs with “many-to-one” and “many-to-many” relationships in order to determine similarities and differences in evolutionary trajectories for the two coral species under investigation, as differentiation in function are suggested by increases and decreases in gene family sizes.

This analysis revealed several striking features. First, among the shared enriched processes for this category of orthologs, we found many processes directly related to the cnidarian-dinoflagellate symbiosis. This partially resembles the results from the one-to-one ortholog analysis with the important difference that independent expansions of genes that map to common processes indicate that cnidarian-dinoflagellate symbioses are actively being shaped within coral species and that different hosts seem to converge on the same processes, indicating convergent evolution. Second, within these processes, homologs of the same or similar genes are repeatedly being expanded across species, as highlighted by the example of three cases of expansion of NLRC3. This indicates that the same genes are potentially subjected to adaptation within and between coral species, arguing that convergent evolution not only happens on the process level, but also on the protein level. Further evolutionary analysis incorporating more species might provide an avenue to identify genes important to coral host adaptation. Last, even though we find expansions of the same genes between species, the extent of duplication is in some cases extremely uneven.

In particular when considering ortholog groups that play a role in innate immunity, the emerging pattern is that both coral species independently expanded ortholog groups belonging to TLRs, TNFRs, and NLRs. This resembles the analysis of Shinzato, et al. ^23^ that found that the *A. digitifera* repertoire of Toll/TLR-related receptors was substantially more complex and diverse than that of *Nematostella* and is further in line with the extensive expansion of NLRs in the coral as reported by Hamada, et al. ^42^. Also, our results are in line with Baumgarten *et al.* ^26^ and Poole and Weis ^43^ who found that the TLR/ILR protein repertoires of the symbiotic sea anemone *Aiptasia* show close similarity to *N. vectensis* with apparently lineage-specific expansion in *A. digitifera*. From the admittedly limited analysis of two coral genomes, it appears, however, that *A. digitifera* shows a more pronounced tendency to duplicate genes in the ortholog groups that are expanded in both corals (many-to-many) (Supplementary Dataset S6). This pattern was also apparent when considering innate immunity-related ortholog groups that are expanded in only one of both corals (“many-to-one”) (although expansions were similar when considering all ortholog groups) (Supplementary Dataset S7). Hence, it will be interesting to see whether *A. digitifera* (and perhaps other *Acropora* species) represent indeed ‘extreme’ cases of expansion of innate immunity-related genes (even within corals) and whether this might even be a hallmark of coral species from the complex clade. With more coral genomes expected to becoming available soon^9^, this would be a fascinating question to pursue. But the analysis of “many-to-one” ortholog groups also revealed that *A. digitifera* and *S. pistillata* seem to diverge on which innate immunity-related genes are expanded. In *A. digitifera* we find NLRs, whereas in *S. pistillata* we find TLRs and TRAFs to be preferentially expanded. Thus, it will be interesting to determine, whether these evolutionary differences might help to pinpoint groups of genes or individual proteins that determine differential specificity to algal symbionts or can be related to differences in physiology, such as thermal tolerance, stress resilience, symbiont transmission mode, and others ^9^.

Taken together, our analyses corroborate recent comparative genomic analyses that showcase how the proteomic information stored in coral genomes has provided the foundation for adapting to a symbiotic, sessile, and calcifying lifestyle of scleractinian corals ^22,23,44^. In particular, our analyses of the core set of conserved proteins and the set of independently expanded ortholog groups in both species underscore the putative importance of the endosymbiotic relationship in determining evolutionary patterns. At the same time, our results demonstrate that coral genomes can be surprisingly disparate as highlighted by extremely uneven or independent expansions of some ortholog groups. It will be important to determine whether the patterns describe here extend to differences between clades and, most importantly, if they are predictive and relevant to a coral’s ability to respond to environmental change.

## Methods

### Organism and isolation of genomic DNA

Colonies of *S. pistillata*, collected at a depth of 5m in front of the Marine Science Station, Gulf of Aqaba, Jordan ^45^, were transferred and maintained at the Centre Scientifique de Monaco in aquaria supplied with flowing seawater from the Mediterranean Sea (exchange rate: 2% h^-1^) at a salinity of 38.2 PPT, pH 8.1 ± 0.1 under an irradiance of 300 μmol photons m^-2^ s^-1^ at 25 ± 0.5 °C. Corals were fed three times a week with frozen krill and live *Artemia salina* larvae. Based on nuclear ITS and mitochondrial COI, coral colonies were typed to be *S. pistillata* clade 4, which is found throughout the northwest Indian Ocean including the Red Sea, the Persian/Arabian Gulf and Kenya ^46^ (Fig. S1). DNA for sequencing libraries was extracted from *S. pistillata* nubbins using a nuclei isolation approach to minimize contamination with algal symbiont DNA. Briefly, cells from a *S. pistillata* nubbin of about 3 cm were harvested in 50 ml of 0.2 M EDTA solution using a water pick and refrigerated at 4 °C. Extracts were first passed through a 100 μm and subsequently through a 40 μm cell strainer (Falcon, Corning, Tewksbury MA, USA) to eliminate most of the zooxanthellae. Next, extracts were centrifuged at 2,000 g for 10 min at 4 °C. The supernatant was discarded and the resulting pellets were homogenized in lysis buffer (G2) of the Qiagen Genomic DNA isolation kit (Qiagen, Hilden, Germany). DNA was extracted following manufacturer’s instructions using genomic-tip 100/G. DNA concentration was determined by O.D. with Epoch Microplate Spectrophotometer (BioTek, Winooski, VT, USA). A check for potential co-isolation of *Symbiodinium* DNA was assessed via PCR targeting the multicopy gene RuBisCO (Genbank accession number AY996050) and did not yield any amplification.

### Genome size estimation

To assess genome size and validate the bioinformatically estimated genome size, we performed a physical measurement of nuclei DNA content using chicken red blood cells (CRBC) as a reference (DNA QC Particles kit, BD Biosciences, San Jose, CA, USA). Extraction and staining of nuclei were performed following the ‘CyStain PI absolute T’ kit (PARTEC #05-5023, Partec, Muenster, Germany) following the manufacturer’s recommendation. Briefly, cells from *S. pistillata* from a nubbin of about 3 cm were harvested using a Water Pick in 50 ml of 0.2 M EDTA solution refrigerated at 4 °C and centrifuged at 2,000 g for 10 min at 4 °C. The cell pellet was resuspended in nuclei extraction buffer, incubated for 15 min at 22 °C and subsequently filtered through a 40 μm cell strainer. Cell lysates were stained with propidium iodide for 60 min at 22 °C, protected from light. Fluorescence signals from nuclear suspensions of separate (i.e., *S. pistillata* or CRBC) and mixed nuclei (i.e., *S. pistillata* and CRBC) were measured on a LSRII Fortessa (BD Biosciences, San Jose, CA, USA) using a 561 nm laser and BP605/40 filter. Based on the known diploid DNA content of chicken erythrocytes of 2.33 pg per cell), coral genome size calculation was determined as follows: sample genome size [pg] = 1.165 x / y (x: fluorescence intensity of your unknown sample; y: fluorescence intensity of CRBC). After calculating mean DNA content per copy of genetic information (1C), genome size can be determined by considering that 1 pg DNA equals 978 Mb. The measurements yielded an estimated *Stylophora pistillata* haploid genome size of 434 Mb (Fig. S2).

### Genome sequencing and assembly

Sequencing libraries were prepared using the Illumina TruSeq DNA kits for paired-end or mate-pair libraries respectively according to the manufacturer’s instructions. A total of 5 paired-end and 8 mate-pair libraries were generated and sequenced on the Illumina HiSeq platform at the KAUST Bioscience Core Facility with exception of the library “miseq”, which was sequenced on the Illumina MiSeq platform (Table S4). An additional mate-pair library (mp05) was generated and sequenced at GATC Biotech (Konstanz, Germany) (Table S4). All data were uploaded to NCBI and are available under Bioproject ID PRJNA281535 (https://www.ncbi.nlm.nih.gov/bioproject/PRJNA281535/).

All sequencing libraries (435x coverage) were trimmed using Trimmomatic version 0.32 ^47^ to remove adaptors, primers, and low quality bases at the ends of sequence reads. Putative PCR duplicates were removed using FastUniq version 1.1 ^48^ to compact the dataset for higher assembly performance. Three-pass digital normalization was performed on all paired-end libraries to reduce data redundancy with khmer ^49^ version 0.7.1 (k=20 C=20, then k=20 C=10). Four paired-end libraries (221x coverage) and four mate-pair libraries (96x coverage) were *de novo* assembled with ALLPATHS-LG release 48961 ^50^ using parameter HAPLOIDIFY=True, and transcriptome data was used to scaffold the assembly with L_RNA_Scaffolder ^51^. All Illumina sequencing libraries were used for scaffolding and gap filling using SSPACE version 1.2 ^52^ and GapFiller version 1.11 ^53^ iteratively for 3 rounds. The above-described procedure yielded an assembly of 358,078,850 bp total contig size and 400,108,361 bp total scaffold size with respective N50s of 24,388 bp and 457,453 bp. Basic genome statistics for contigs and scaffolds were generated using the perl script (http://korflab.ucdavis.edu/datasets/Assemblathon/Assemblathon2/Basic_metrics/assemblathon_stats.pl) used to validate assemblies in the “Assemblathon 2 Contest” ^54^. The estimated genome size as per ALLPATHS-LG was reported at 433 Mb, and the assembled contig and scaffold lengths provide a genome coverage of ~83% and 92%, respectively. Further information regarding genome statistics are provided in Table 1 and Table S1.

### Identification and removal of contaminating sequences

To identify and remove sequences likely to have originated from dinoflagellate, bacterial, or viral contaminants, a custom script was written employing the following strategy. BLASTN searches were conducted against six databases: the genomes of *S. minutum* ^55^ and *S. microadriaticum* ^56^; complete bacterial genomes (ftp://ftp.ncbi.nih.gov/genomes/Bacteria/all.fna.tar.gz), draft bacterial genomes (ftp://ftp.ncbi.nih.gov/genomes/Bacteria_DRAFT/), and complete viral genomes (ftp://ftp.ncbi.nih.gov/genomes/Viruses/all.fna.tar.gz) databases from NCBI; and the viral database PhAnToMe (http://phantome.org). All databases were retrieved in July 2014. As the lengths of the query and hit sequences were up to hundreds of kilobases, a combination of cutoffs (total bit score > 1,000, e-value ≤ 10^-20^) was used to identify scaffolds with significant sequence similarities to non-coral sequences representing potential contaminants. This procedure yielded 41 scaffolds that displayed significant similarity in over 50% of their non-N sequences, and thus were considered to have originated from bacterial contaminants and removed from the final assembly.

### Annotation of repetitive elements

*De novo* identification of species-specific repeat regions in the genome assembly of *S. pistillata* was performed using RepeatScout (version 1.0.5) ^57^ with an l-mer size of l = 16 bp. Using the default settings, 10,224 distinct repeat motifs were identified that occurred ≥10 times. Annotation of these repeats was performed as described previously ^26^ using three different methods: (i) RepeatMasker (version 4.0.2) ^58^ using RepBase version 19.07, (ii) TBLASTX against RepBase version 19.07, and (iii) BLASTX against a custom-made non-redundant database of proteins encoded by transposable elements (TEs; NCBI keywords: retrotransposon, transposase, reverse transcriptase, gypsy, copia). The best annotation among the three methods was chosen based on alignment coverage and score. The repeat motifs identified in this way and the set of known eukaryotic TEs from RepBase (May 2014 release) were then used to locate and annotate the repeat elements in the assembled genome using RepeatMasker (version 4.0.2). The repeat identification and annotation for the *A. digitifera* genome assembly was performed *sensu* Baumgarten, et al.^26^ (Table S5).

### Reference transcriptome sequencing and assembly

Total RNA was extracted from *S. pistillata* nubbins subjected to different pH treatments. Briefly, nubbins were cultured in triplicates at pH 7.2, 7.6, 7.8, and 8.1 for 24 months prior to extraction. The RNA preparation was performed as described in Liew, et al. ^59^ and strand-specific sequencing libraries were generated using the NEBNext Ultra Directional RNA Library Prep Kit for Illumina (NEB, Ipswitch, MA, USA). A total of 12 libraries were generated and sequenced on 2 lanes of the Illumina HiSeq platform at the KAUST Bioscience Core Facility (KAUST, Thuwal, KSA), producing 924 million read pairs with 900x coverage. Sequence reads from all libraries were trimmed using Trimmomatic version 0.32 to remove adaptors, primers, and low quality read ends (base quality < 30). Further, all reads shorter than 35 bp were removed. PhiX reads were removed using Bowtie2 ^60^ and putative PCR duplicates were removed with PRINSEQ-lite version 0.20.3 ^61^. After that, all libraries were merged and error correction was carried out using ErrorCorrectReads.pl (ALLPATHS-LG). The resulting merged library was assembled *de novo* using Trinity ^62^ release 20140413 with strand-specific parameter (-- SS_lib_type RF --min_kmer_cov 5 --normalize_reads) yielding a total of 89,208 assembled transcripts. The reference transcriptome is available at reefgenomics.org ^63^ at http://spis.reefgenomics.org.

### Gene model prediction

All 89,208 transcripts from the reference transcriptome (see above) were mapped to the genome assembly and filtered by PASA release 20140417 ^62^ to create a training set for AUGUSTUS version 3.0.2 ^64,65^. The training set was filtered using the following steps: (1) incomplete transcripts were removed, (2) transcripts with less than 3 exons were removed, (3) transcripts with ambiguous 5’ or 3’ untranslated regions (UTRs) were removed, (4) redundant transcripts were removed as indicated by BLASTP, (5) transcripts harboring repeat sequences were removed based on BLASTN against the repeat library generated by RepeatScout (see above). This yielded 2,844 transcripts and corresponding mapping information that were used to train AUGUSTUS. To improve prediction accuracy, a ‘hints’ file indicating the locations of matching transcripts was generated by mapping all 89,208 transcripts from the reference transcriptome to the genome assembly using BLAT and the AUGUSTUS script blat2hints.pl. Using the ‘hints’ file, AUGUSTUS was used for *ab initio* prediction of gene models from the genome assembly and PASA was used subsequently for comparison and completion of the gene models.

### Genome protein set completeness analysis

Completeness of the *S. pistillata* genome was assessed using the CEGMA (Core Eukaryotic Genes Mapping Approach) pipeline ^66^ that is based on a set of core eukaryotic genes (CEGs) from six model organisms (*Homo sapiens, Drosophila melanogaster, Arabidopsis thaliana, Caenorhabditis elegans, Saccharomyces pombe*, and *Saccharomyces cerevisiae*). CEGMA searches for existence of 248 highly conserved genes in the genomic protein set using an approach based on BLASTP and subsequent validation using Hidden Markov Models generated for the core gene set in order to estimate the completeness of a given genomic gene set.

### Coral genomic protein set composition

BLASTP searches of gene models from *S. pistillata* and *A. digitifera* were performed against two different databases. Using *S. pistillata* to illustrate the procedure, two databases were created: a ‘non-coral’ database, and a ‘with-coral’ database. The former consisted of *Aiptasia* gene models ^26^ and the NCBI ‘nr’ database (Nov. 2015 release); the latter included gene models from *A. digitifera*, in addition to the sequences from the ‘non-coral’ database. Species names contained within annotations for the best hits (e-value ≤ 10^-5^) were parsed and fed into a python script that obtained the full taxonomic hierarchy for the respective organisms via an API hosted by Encyclopedia of Life (http://eol.org/api) ^67^. Based on the resulting hierarchies, best hits were grouped based on their respective genus. Chord diagrams were drawn using Circos ^68^. For visual clarity, only the six most frequent genera were shown and all others were collapsed into “Others”.

### Protein set annotation

The final set of predicted proteins derived their annotations from UniProt (i.e., SwissProt and TrEMBL) or the NCBI ‘nr’ databases (Table S1, Supplementary Dataset S8), similar to the pipeline described previously ^26,59^. Briefly, genomic protein models were subjected to a BLASTP search against SwissProt and TrEMBL databases (June 2014 release). GO terms associated with SwissProt and TrEMBL hits were obtained from UniProt-GOA (July 2014 release) ^69^. If the best-scoring hit of the BLASTP search did not yield any GO annotation, further hits (up to 20 hits, e-value ≤ 10^-5^) were considered, and the best-scoring hit with available GO annotation was used. If none of the SwissProt hits had GO terms associated with them, the TrEMBL hits were processed similarly. Using this procedure, 21,446 genes (83.2% of the 25,769 gene models) were annotated and had at least one GO term associated with them (17,506 proteins had GO annotations via Swiss-Prot, while the remaining 3,940 were from TrEMBL) (Table S1). A majority of the annotations were based on strong alignments to existing sequences within the SwissProt and TrEMBL databases: 19,060 genes had e-values ≤ 10^-10^, 15,637 of these had e-values ≤ 10^-20^. Proteins that had no matches to either database were subjected to an additional search against the NCBI ‘nr’ database (e-value ≤ 10^-5^). An additional 1,466 proteins were annotated this way. A small fraction of proteins (2,857, 11.1%) had no hits to any of the three databases. A similar procedure was performed for the *A. digitifera* gene models to eliminate potential biases stemming from the use of different annotation pipelines (Supplementary Dataset S9).

### Ortholog identification, category assignment, and GO enrichment analyses

Orthologs between *S. pistillata* (n = 25,679 genes) and the *A. digitifera* V1 gene set (n = 23,523 genes) (http://marinegenomics.oist.jp/genomes/downloads?project_id=3) were identified using Inparanoid v4.1 ^70^ and assigned to four categories according to their evolutionary relationships: (i) one-to-one orthologs, (ii) many-to-one and many-to-many orthologs, (iii) proteins without easily discernible orthologous relationships, and (iv) species-specific proteins without homologs in other species. Gene Ontology (GO) enrichment analyses of orthologs from all categories were conducted by testing annotations from genes belong to a given category relative to annotations from all genes of that species. For instance, in order to investigate enrichment of biological functions of the one-to-one orthologs, GO terms within the list of 6,302 *S. pistillata* genes were tested for enrichment relative to all *S. pistillata* genes with at least one annotated GO term (21,446 genes). Similarly, the 6,302 *A. digitifera* genes were tested against *A. digitifera* genes with at least one annotated GO term (18,544 genes). GO enrichment analyses were conducted using topGO (v2.24.0) ^71^ with the “weight01” settings. The threshold for significance was *p* < 0.05. The *p* values were not corrected for multiple testing as non-independent tests were carried out on each GO term.

### Phylogenetic trees

Sequences from ortholog groups of interest were first aligned using MUSCLE ^72^, and aligned sequences were then trimmed with trimAl v1.4.1 ^73^ with the “-automated1” flag. The alignments were subsequently constructed using RAxML v8.2.9 ^74^ with 1,000 bootstraps (-m PROTGAMMAJTT -x 12345 -p 12345 -N 1000 -f a). Trees were viewed and exported to a graphical format using FigTree v1.4.2 (http://tree.bio.ed.ac.uk/software/figtree/).

## Acknowledgements

We would like to thank the Bioscience Core Lab at KAUST for sequencing. Further, we would like to thank Adrian Carr and Gos Micklem for support with an initial genome assembly. Research reported in this publication was supported by King Abdullah University of Science and Technology (KAUST).

## Author contributions

CRV, MA, YL, YJL, SB designed and conceived the study; DZ, ST, DA generated data; YJL, SB, YL, MA, CRV analyzed data; MA, DZ, ST, DA, CRV contributed reagents/tools/materials; CRV wrote the manuscript with contributions from MA, YJL, YL, and SB; all authors read and approved the final manuscript.

## Additional Information

### Accession codes

The genome assembly, gene models, and protein models described in this study are available for download at http://spis.reefgenomics.org/download. A JBrowse genome browser is available at http://spis.reefgenomics.org/jbrowse. A BLAST server for the *Stylophora pistillata* genome is available at http://spis.reefgenomics.org/blast/. Raw sequence data reported are deposited at NCBI under the accession number PRJNA281535 (https://www.ncbi.nlm.nih.gov/bioproject/PRJNA281535/).

## References

1 Porter, J. W. & Tougas, J. I. in Encyclopedia of Biodiversity (ed Simon Asher Levin) 73–95 (Elsevier, 2001).

2 Wilkinson, C. e. Status of Coral Reefs of the World: 2008. Global Coral Reef Monitoring Network and Reef and Rainforest Research Center, Townsville, Australia (2008).

3 Hughes, T. P. et al. Climate change, human impacts, and the resilience of coral reefs. Science 301, 929–933 (2003).

4 Hoegh-Guldberg, O. et al. Coral reefs under rapid climate change and ocean acidification. Science 318, 1737–1742 (2007).

5 Maynard, J. et al. Projections of climate conditions that increase coral disease susceptibility and pathogen abundance and virulence. Nature Clim. Change 5, 688–694 (2015).

6 Carpenter, K. E. et al. One-third of reef-building corals face elevated extinction risk from climate change and local impacts. Science 321, 560–563 (2008).

7 Forest, R., Victor, S., Farooq, A. & Nancy, K. Diversity and distribution of coral-associated bacteria. Marine Ecology Progress Series 243, 1–10 (2002).

8 Knowlton, N. & Rohwer, F. Multispecies Microbial Mutualisms on Coral Reefs: The Host as a Habitat. The American Naturalist 162, S51–S62 (2003).

9 Voolstra, C. et al. The ReFuGe 2020 Consortium—using “omics” approaches to explore the adaptability and resilience of coral holobionts to environmental change. Frontiers in Marine Science 2, 68 (2015).

10 Daniels, C. et al. Metatranscriptome analysis of the reef-buidling coral Orbicella faveolata indicates holobiont response to coral disease. Frontiers in Marine Science 2 (2015).

11 Theis, K. R. et al. Getting the hologenome concept right: An eco-evolutionary framework for hosts and their microbiomes. bioRxiv (2016).

12 McFall-Ngai, M. et al. Animals in a bacterial world, a new imperative for the life sciences. Proceedings of the National Academy of Sciences 110, 3229–3236 (2013).

13 Rosenberg, E., Koren, O., Reshef, L., Efrony, R. & Zilber-Rosenberg, I. The role of microorganisms in coral health, disease and evolution. Nature Reviews: Microbiology 5, 355–362 (2007).

14 Bay, R. A. & Palumbi, S. R. Rapid Acclimation Ability Mediated by Transcriptome Changes in Reef-Building Corals. Genome biology and evolution 7, 1602–1612 (2015).

15 Seneca, F. O. & Palumbi, S. R. The role of transcriptome resilience in resistance of corals to bleaching. Molecular ecology 24, 1467–1484 (2015).

16 Barshis, D. J. et al. Genomic basis for coral resilience to climate change. Proceedings of the National Academy of Sciences of the United States of America 110, 1387–1392 (2013).

17 Kenkel, C. D., Meyer, E. & Matz, M. V. Gene expression under chronic heat stress in populations of the mustard hill coral (Porites astreoides) from different thermal environments. Molecular ecology 22, 4322–4334 (2013).

18 Hume, B. C. C. et al. Ancestral genetic diversity associated with the rapid spread of stress-tolerant coral symbionts in response to Holocene climate change. Proceedings of the National Academy of Sciences 113, 4416–4421 (2016).

19 Ziegler, M. et al. Coral microbial community dynamics in response to anthropogenic impacts near a major city in the central Red Sea. Marine pollution bulletin 105, 629–640 (2016).

20 Ziegler, M. et al. Biogeography and molecular diversity of coral symbionts in the genus Symbiodinium around the Arabian Peninsula. Journal of Biogeography Accepted (2016).

21 Ziegler, M., Seneca, F. O., Yum, L. K., Palumbi, S. R. & Voolstra, C. R. Bacterial community dynamics are linked to patterns of coral heat tolerance. Nature Communications Accepted (2016).

22 Bhattacharya, D. et al. Comparative genomics explains the evolutionary success of reef-forming corals. eLife 5, e13288 (2016).

23 Shinzato, C. et al. Using the Acropora digitifera genome to understand coral responses to environmental change. Nature 476, 320–323 (2011).

24 Putnam, N. H. et al. Sea anemone genome reveals ancestral eumetazoan gene repertoire and genomic organization. Science 317, 86–94 (2007).

25 Chapman, J. A. et al. The dynamic genome of Hydra. Nature 464, 592–596 (2010).

26 Baumgarten, S. et al. The genome of Aiptasia, a sea anemone model for coral symbiosis. Proceedings of the National Academy of Sciences 112, 11893–11898 (2015).

27 Park, E. et al. Estimation of divergence times in cnidarian evolution based on mitochondrial protein-coding genes and the fossil record. Molecular Phylogenetics and Evolution 62, 329–345 (2012).

28 Simpson, C., Kiessling, W., Mewis, H., Baron-Szabo, R. C. & Müller, J. Evoluationary Diversification of Reef Corals: A comparison of the molecular and fossil records. Evolution 65, 3274–3284 (2011).

29 Khalturin, K., Hemmrich, G., Fraune, S., Augustin, R. & Bosch, T. C. G. More than just orphans: are taxonomically-restricted genes important in evolution? Trends in Genetics 25, 404–413 (2009).

30 Zdobnov, E. M. et al. Genome and Proteome Analysis of Anopheles gambiae and Drosophila melanogaster. Science 298, 149–159 (2002).

31 Parra, G., Bradnam, K. & Korf, I. CEGMA: a pipeline to accurately annotate core genes in eukaryotic genomes. Bioinformatics 23, 1061–1067 (2007).

32 Gaunt, M. W. & Miles, M. A. An Insect Molecular Clock Dates the Origin of the Insects and Accords with Palaeontological and Biogeographic Landmarks. Molecular Biology and Evolution 19, 748–761 (2002).

33 Voolstra, C. R. et al. Rapid Evolution of Coral Proteins Responsible for Interaction with the Environment. PLoS ONE 6, e20392 (2011).

34 Aparicio, S. et al. Whole-Genome Shotgun Assembly and Analysis of the Genome of Fugu rubripes. Science 297, 1301–1310 (2002).

35 Rädecker, N., Pogoreutz, C., Voolstra, C. R., Wiedenmann, J. & Wild, C. Nitrogen cycling in corals: the key to understanding holobiont functioning? Trends in Microbiology 23, 490–497 (2015).

36 Ganot, P. et al. Structural Molecular Components of Septate Junctions in Cnidarians Point to the Origin of Epithelial Junctions in Eukaryotes. Molecular Biology and Evolution 32, 44–62 (2015).

37 Kitahara, M. V., Cairns, S. D., Stolarski, J., Blair, D. & Miller, D. J. A Comprehensive Phylogenetic Analysis of the Scleractinia (Cnidaria, Anthozoa) Based on Mitochondrial CO1 Sequence Data. PLoS ONE 5, e11490 (2010).

38 Shearer, T. L., Van Oppen, M. J., Romano, S. L. & Worheide, G. Slow mitochondrial DNA sequence evolution in the Anthozoa (Cnidaria). Molecular ecology 11, 2475–2487 (2002).

39 Kitahara, M. V. et al. The “Naked Coral” Hypothesis Revisited – Evidence for and Against Scleractinian Monophyly. PLoS ONE 9, e94774 (2014).

40 Fukami, H. et al. Mitochondrial and Nuclear Genes Suggest that Stony Corals Are Monophyletic but Most Families of Stony Corals Are Not (Order Scleractinia, Class Anthozoa, Phylum Cnidaria). PLoS ONE 3, e3222 (2008).

41 Wells, J. in Treatise on Invertebrate Paleontology. Part F. Coelenterata (ed RC Moore) 328–440 (Geological Society of America & University of Kansas Press, 1956).

42 Hamada, M. et al. The Complex NOD-Like Receptor Repertoire of the Coral Acropora digitifera Includes Novel Domain Combinations. Molecular Biology and Evolution 30, 167–176 (2013).

43 Poole, A. Z. & Weis, V. M. TIR-domain-containing protein repertoire of nine anthozoan species reveals coral-specific expansions and uncharacterized proteins. Developmental and comparative immunology 46, 480–488 (2014).

44 Zoccola, D. et al. Bicarbonate transporters in corals point towards a key step in the evolution of cnidarian calcification. Scientific Reports 5, 9983 (2015).

45 Tambutte et al. A compartmental approach to the mechanism of calcification in hermatypic corals. The Journal of experimental biology 199, 1029–1041 (1996).

46 Keshavmurthy, S. et al. DNA barcoding reveals the coral “laboratory-rat", Stylophora pistillata encompasses multiple identities. Scientific Reports 3, 1520 (2013).

47 Bolger, A. M., Lohse, M. & Usadel, B. Trimmomatic: a flexible trimmer for Illumina sequence data. Bioinformatics 30, 2114–2120 (2014).

48 Xu, H. et al. FastUniq: A Fast De Novo Duplicates Removal Tool for Paired Short Reads. PLoS ONE 7, e52249 (2012).

49 Crusoe, M. R. et al. The khmer software package: enabling efficient nucleotide sequence analysis. F1000Research 4, 900 (2015).

50 Gnerre, S. et al. High-quality draft assemblies of mammalian genomes from massively parallel sequence data. Proceedings of the National Academy of Sciences 108, 1513–1518 (2011).

51 Xue, W. et al. L_RNA_scaffolder: scaffolding genomes with transcripts. BMC Genomics 14, 604 (2013).

52 Boetzer, M., Henkel, C. V., Jansen, H. J., Butler, D. & Pirovano, W. Scaffolding pre-assembled contigs using SSPACE. Bioinformatics 27, 578–579 (2011).

53 Boetzer, M. & Pirovano, W. Toward almost closed genomes with GapFiller. Genome Biology 13, 1–9 (2012).

54 Bradnam, K. R. et al. Assemblathon 2: evaluating de novo methods of genome assembly in three vertebrate species. GigaScience 2, 10–10 (2013).

55 Shoguchi, E. et al. Draft assembly of the Symbiodinium minutum nuclear genome reveals dinoflagellate gene structure. Current biology: CB 23, 1399–1408 (2013).

56 Aranda, M. et al. Genome analysis of coral dinoflagellate symbionts highlights evolutionary adaptations to a symbiotic lifestyle. Scientific Reports Accepted (2016).

57 Price, A. L., Jones, N. C. & Pevzner, P. A. De novo identification of repeat families in large genomes. Bioinformatics 21, i351–i358 (2005).

58 Smit, A., Hubley, R. & Green, P. RepeatMasker Open-4.0., http://www.repeatmasker.org. (2013-2015).

59 Liew, Y. J. et al. Identification of microRNAs in the coral Stylophora pistillata. Plos One 9 (2014).

60 Langmead, B. & Salzberg, S. L. Fast gapped-read alignment with Bowtie 2. Nature methods 9, 357–359 (2012).

61 Schmieder, R. & Edwards, R. Quality control and preprocessing of metagenomic datasets. Bioinformatics 27, 863–864 (2011).

62 Haas, B. J. et al. De novo transcript sequence reconstruction from RNA-Seq: reference generation and analysis with Trinity. Nature protocols 8, 10.1038/nprot.2013.1084 (2013).

63 Liew Y.J., Aranda M. & C.R., V. reefgenomics.org—a repository for marine genomics data. Database Accepted (2016).

64 Stanke, M. et al. AUGUSTUS: ab initio prediction of alternative transcripts. Nucleic acids research 34, W435–439 (2006).

65 Stanke, M. & Waack, S. Gene prediction with a hidden Markov model and a new intron submodel. Bioinformatics 19 Suppl 2, ii215–225 (2003).

66 Parra, G., Bradnam, K., Ning, Z., Keane, T. & Korf, I. Assessing the gene space in draft genomes. Nucleic acids research 37, 289–297 (2009).

67 Parr, C. S. et al. The Encyclopedia of Life v2: Providing Global Access to Knowledge About Life on Earth. Biodiversity Data Journal, e1079 (2014).

68 Krzywinski, M. et al. Circos: An information aesthetic for comparative genomics. Genome Research 19, 1639–1645 (2009).

69 Dimmer, E. et al. The UniProt-GO Annotation database in 2011. Nucleic acids research 40, D565 - D570 (2012).

70 Sonnhammer, E. L. L. & Östlund, G. InParanoid 8: orthology analysis between 273 proteomes, mostly eukaryotic. Nucleic acids research (2014).

71 Alexa, A. & Rahnenfuhrer, J. topGO: enrichment analysis for gene ontology. R package version 2.8 (2010).

72 Edgar, R. C. MUSCLE: multiple sequence alignment with high accuracy and high throughput. Nucleic acids research 32, 1792–1797 (2004).

73 Capella-Gutierrez, S., Silla-Martinez, J. M. & Gabaldon, T. trimAl: a tool for automated alignment trimming in large-scale phylogenetic analyses. Bioinformatics 25, 1972–1973 (2009).

74 Stamatakis, A. RAxML version 8: a tool for phylogenetic analysis and post-analysis of large phylogenies. Bioinformatics 30, 1312–1313 (2014).

